# Cross-Neutralizing Monoclonal Antibodies with Broad Activity Against Human and Bat-Derived SARS-Related Coronaviruses

**DOI:** 10.1101/2025.08.14.670296

**Authors:** Martin Mayora Neto, Kelly da Costa, Diego Cantoni, Mariliza Derveni, Edward Wright, Heinz Hoschuetzky, Franz Kaufmann, Petra Schuessler, Nigel Temperton

**Affiliations:** Viral Pseudotype Unit, Medway School of Pharmacy, University of Kent, Chatham, United Kingdom; Viral Pseudotype Unit, University of Sussex, Brighton, United Kingdom; Nanotools Antikorpertechnik GmbH & Co. KG Tscheulinstraße 21, Teningen, Germany

## Abstract

Effective therapeutics for severe cases of SARS-CoV-2 are still needed. As new variants of concern emerged an increase in hospitalizations was observed, especially in non-vaccinated individuals, immunocompromised individuals and the elderly, whereby treatment options became challenging. Several monoclonal antibodies (mAb) are being evaluated for approval due to their ability to neutralize the virus. Here, we report monoclonal antibodies targeting the Spike (S) with a strong neutralization profile against lentiviral and VSV pseudotypes displaying the Spikes of SARS-CoV-1, SARS-CoV-2 and variants of concern, in addition to two bat coronaviruses. We found several mAbs that were able to bind and neutralize a broad set of variants, with one mAb able to neutralize SARS-CoV-1, SARS-CoV-2, RaTG13 and WIV16. Their binding affinities were also characterized, and several mAbs were in the picomolar range against different variants.

These results indicate that their use as a cocktail of monoclonal antibodies to treat patients infected with variants associated with severe disease and immune escape can be explored as an option. Furthermore, the cross reactivity observed by one mAb may reveal further insight into vulnerable portions of the Spike protein, which may become valuable future targets.

## Introduction

Zoonotic transmission of viruses has resulted in multiple outbreaks with pandemic potential^1^. Coronaviruses have been responsible for concerning outbreaks and pandemics since SARS-CoV in 2003^2,3^, MERS-CoV in 2012^4^, and most recently the ongoing pandemic of SARS-CoV-2^5,6^.

Since the emergence of SARS-CoV-2, the aetiological agent of COVID-19, several effective vaccines are on the market as a result of collaborative global efforts^7^. The adenovirus and mRNA platforms have been very successful in preventing severe disease^7^. However, emerging variants of concern (VOC) have been associated with increased transmission and vaccine breakthrough infections leading to an increase in hospitalizations during several waves^8–10^. The Alpha variant emerged in south-east England in 2020 exhibiting increased transmission and accounting for the majority of cases in Europe by April 2021^11^. Concomitantly the Beta variant emerged in South Africa, also with increased transmissibility and immune evasion^12,13^. The Delta variant was first identified in India and quickly overtook Alpha and Beta variants, of which the latter exhibited strong immune evasion properties due to key mutations in the receptor binding domain. More recently, the Omicron BA.1 variant rapidly displaced Delta, due to its increased virulence and strong immune evasion^13^. Omicron BA.1 was unique in that it contained the most mutations in the Spike observed when it emerged during the pandemic and also exhibited a switch in entry mechanism, preferring the endosomal entry route^14–16^. Omicron and its multiple iterations has now dominated cases around the world, with BA.5 and more recently BA2.86, BQ.1.1 and XBB.1, highlighting its high replication dynamics coupled with strong immune evasion ^17–19^.

The Spike protein on the surface of the virion is the major determinant of tropism^20^. It interacts with its cellular receptor promoting entry. Coronaviruses utilize a range of receptors including angiotensin-converting enzyme (ACE2) for SARS-CoV, SARS-CoV-2 and NL63; dipeptidyl peptidase-4 (DPP4) for MERS-CoV; human aminopeptidase N (hAPN) for 229E; and sialoglycan-based receptors containing 9-*O*-acetylated sialic acid (9-*O*-Ac-Sia) for HKU1 and OC43^21^. The S protein is the main target of neutralizing antibodies, which prevent viral entry. As a result, most vaccination platforms have used S as their immunogen, with varying degrees of success^22,23^.

Monoclonal antibodies are being evaluated as therapeutic agents for patients with severe COVID-19 or those who generate suboptimal immune responses, such as immunocompromised individuals^24–27^. Some of the monoclonal antibodies being evaluated in clinical trials have raised the concern of reduced neutralization of variants Beta (B.1.351) and Alpha (B.1.1.7) for instance^28^, therefore a cocktail of mAbs targeting different epitopes would be less likely to result in mutational escape and antibody resistance^29^. Furthermore, mAbs with cross-neutralizing potential against other (related) viruses such as SARS-CoV might be useful in future outbreaks^30^, resulting in a global effort to identify pan-sarbecovirus monoclonal antibodies^31–33^. This can be achieved as several studies have shown presence of cross-reactive antibodies generated by SARS-CoV-2 targeted vaccination against RaTG13^34^, SARS-CoV-1^35^ and WIV16^36,37^.

The monoclonal antibody cocktail Evusheld™ (tixagevimab, cilgavimab) was approved for emergency use as prophylaxis as well as treatment in certain patients with severe disease^38^. However, since the emergence of Omicron variants, it lost its effectiveness^38^, resulting in the FDA withdrawing authorization for its use in the US in 2023.

The need for high containment facilities for working with wild-type SARS-CoV-2 creates an R&D bottleneck. Pseudotype viruses (PV) have been crucial to circumvent this issue as they are non-replicative viruses containing a reporter gene, displaying S on their surface that can be handled in low containment^39–44^. Two pseudotype platforms have been used in this comparative study. Vesicular stomatitis virus (VSV) pseudotypes harbour an almost complete genome, where a luciferase reporter gene replaces the G gene. Recombinant virions have to be generated and amplified by providing the G cell surface protein *in trans*^45^. These are then used to infect cells previously transfected with a coronavirus S protein, resulting in a PV with a VSV core displaying S and capable of a single round of infection.

Lentiviral PVs are generated through a 3-plasmid transfection method including S, a lentiviral vector encoding a luciferase reporter gene, and a plasmid encoding a HIV-1 core including all the enzymes needed for maturation, and integration of the reporter gene into the target cell genome^46,47^. They can be used in pseudotype neutralization assays (pMN) resulting in a reduction in reporter gene expression if neutralizing antibodies prevent PV entry^48^. These platforms act as an alternative to wild-type virus neutralization assays allowing for relatively rapid results, and permitting high-throughput screens of potential neutralizing agents against viruses^49^.

In this study we assessed both VSV and lentiviral PV platforms bearing the Spikes of SARS-CoV-1, SARS-CoV-2, previous VOCs, and two bat coronaviruses, RaTG13 and WIV16 in a pMN, against a panel of monoclonal antibodies derived from immunized mice. We also characterized their binding affinities and neutralization profile with a Luminex assay. Their neutralization was additionally evaluated in an ELISA-based surrogate neutralization test (cPASS) in order to assess whether other strategies that do not require tissue culture may be used as a sensitive and rapid alternative to neutralization assays.

## Materials And Methods

### Cell lines and plasmids

HEK293T/17 (ATCC CRL-11268), BHK21 and Huh-7 cells were maintained in Dulbecco Modified Eagle Medium (DMEM with: 4.5 g/l Glucose, stab. Glutamine, Sodium pyruvate, 3.7 g/l NaHCO3, Pan Biotech UK Ltd P04-04510) with 10% v/v FBS and 100 U/mL penicillin/100 µg/mL streptomycin (Pan Biotech).

CHO-K1 cells were a gift from Dr Giada Mattiuzzo, NIBSC, Potters Bar and maintained in Ham F12 medium (Pan Biotech) with 10% FBS and 100 U/mL penicillin/100 µg/mL streptomycin (Pan Biotech).

Spike SARS-CoV-2 (Wuhan) and Delta (B.1.617.2) plasmids have been described previously^50^. Alpha (B.1.1.7)-pI.18 (#101023), Beta (B.1.351)-pI.18 (#101024) and SARS2 (D614G)-pCAGGS (#100985) were gifts from the Medicines & Healthcare Products Regulatory Agency.

Omicron plasmids BA.1, BA.4/5, BA2.75 (pcDNA3.1) were gifts from G2P. BA2.86-pcDNA3.1 was a gift from Dr Joe Grove (MRC-University of Glasgow Centre for Virus Research). MERS^47^, VSV-G^51^ and WIV16^52^ plasmids have been described previously. MT126 pRRL-SFFV-ACE2-IRES puromycin (Addgene plasmid #145839) and MT131 pRRL-SFFV-TMPRSS2.v1-IRES hygromycin were gifts from Dr Caroline Goujon^53^.

### Monoclonal antibodies and Luminex assay

For the generation of monoclonal antibodies, NanoTools’ proprietary high responder mice (NT-HRM) were immunized with particle-based vaccines generated with Spike protein / RBD of Wu SARS-CoV-2. PEG-mediated cell fusions were performed according to Köhler & Milstein using PAI myeloma as fusion partner^54^. Hit selection was performed by Luminex assays for target specificity (high affinity in the low pM range), blocking of the ACE2-Spike trimer interaction, and cross-reactivity between SARS-CoV2 wt and VOC.

Monoclonal Antibodies were purified from serum-free supernatants from subcloned hybridoma cell lines by thiophilic adsorption chromatography. Antibody affinity (KD) and neutralization (IC_50_) were determined by Luminex-based assays.

Analyte proteins were conjugated to COOH activated Luminex beads via carbodiimide chemistry. Biotinylated analyte proteins were bound to streptavidin coated Luminex beads. Binding assays were performed by incubation of Nanotools antibodies with target-loaded beads until equilibrium was reached. Bound antibody was detected with biotinylated anti-mouse antibodies followed by fluorochrome labelled streptavidin. In order to determine dissociation constants (Kd) binding assays were performed using a dilution series of antibodies (1-20000 pM). When fluorescent signals (MFI) are plotted against antibody concentration, the concentration at midpoint MFI corresponds to the Kd value.

*In vitro* assays assessing the neutralization of ACE2-Fc binding to immobilized RBD protein were implemented by incubating Nanotools antibodies with target-loaded beads until equilibrium was reached. After incubation with soluble ACE2-Fc (600 pM) the extent of ACE2-Fc binding was determined by phycoerythrin-labelled anti-human Fc antibody. IC_50_ values correspond to the antibody concentration at midpoint MFI.

### Vesicular Stomatitis Virus pseudotypes

Recombinant VSV (rVSVΔG), in which the glycoprotein (G) gene was replaced with the firefly luciferase gene, was generated and recovered from BHK21 cells (Kerafast Inc.) adapted from a previously optimised protocol^55^. Briefly, BHK21 cells were infected with the recombinant Fowlpox helper virus (FPV-T7) at an MOI of 3 in 6-well plates. The inoculum was removed after 2h, cells washed and an optiMEM/DNA/TransIT2020 transfection mix (500 µL) containing support plasmids NϕT (3 µg/well), PϕT (5 µg/well), GϕT (8 µg/well), LϕT (1 µg/well), pVSVΔG-luc (5 µg/well) was added and incubated for 2h at 37°C before 1.5 mL of DMEM (supplemented with 5% FBS) was added. The following day the medium was replaced to remove the transfection mix. rVSVΔG was harvested 48h after transfection and filtered through a 0.45 µm filter. Stocks were subjected to infectivity assays and TCID_50_ calculations. Amplification of rVSVΔG stocks was done by infection of pre-transfected BHK21 cells with 2 µg of VSV-G at an MOI 0.1. Once rVSVΔG stocks were amplified, VSV PVs were generated on HEK293T/17 producer cells, seeded the day before transfection with (∼5 x 10^6^ cells) in T75 flasks. Next, a DNA mix of 1 µg Spike plasmid was prepared in 200 µL optiMEM (Gibco), 3 µL of FugeneHD (Promega) was added and incubated for 15 min at room temperature. After replacing the media, the transfection mix was added and cells incubated at 37°C 5% CO_2_ overnight. The following day, media was removed and the cells infected with rVSVΔG MOI 1-5 in 4 mL media for 2-3h. The inoculum was discarded, cells washed in DPBS followed by addition of 10 mL complete media. PVs were harvested 24h and 48h post infection, filtered through 0.45 µm cellulose acetate sterile filters and stored at −80°C.

### Lentiviral pseudotypes

Lentiviral pseudotypes were generated as previously described^48,50^. Briefly, 1 x 10^6^ producer HEK293T/17 cells were seeded in a T75 flask the day before transfection. For transfection, a DNA mix was prepared in 200 µL optiMEM containing 1 µg Spike plasmid (pcDNA3.1 or pCAGGS), 1.5 µg pCSFLW luciferase reporter and 1 µg p8.91 HIV-1 gag-pol plasmids. After 5 min, 11 µL FugeneHD was added and incubated for 15 min at room temperature. Next, cell media was replaced with 10 mL complete DMEM and the transfection mix added to the flask. Supernatant was harvested 48-72h post transfection, filtered through a 0.45 µm cellulose acetate filter and stored at −80°C.

### TCID_50_

In a white, flat-bottom, sterile Nunc 96-well microplate, 25 µL of PV supernatant was added in four replicates. Next, 100 µL of complete medium was added to all wells and a 5-fold dilution series was performed. The plate was centrifuged (ELMI CM-6MT centrifuge) for 3 s at 400 rpm (∼21 g) and target cells (5 x 10^4^/well) were added and incubated for 48h at 37°C, 5% CO_2_. After 24h, the media was removed and discarded; Bright-Glo reagent was added to the plate and incubated at room temperature for 5 min before measuring luminescence. The luminescence values were used to calculate the PV titre (TCID_50_/mL) using the Reed-Muensch method^56^. The cumulative number of positive and negative wells for PV infection at each dilution was determined and the percentage calculated for each. The threshold value for a positive well was set at 2.5 x the average luminescence value of the cell only negative controls.

### Pseudotype virus neutralization assay (pMN)

A 2-fold serial dilution of monoclonal antibodies was set up in duplicate in white, flat-bottom, sterile Nunc 96-well microplates. PV was then diluted in complete medium to achieve ∼10^5^ – 10^6^ RLU or ∼10^2^ TCID_50_ per well. A PV only control (0% neutralization) and a cell only control (100% neutralization) were included. The plate was centrifuged for 1 min at 400 rpm (∼21 g) then incubated at 37°C, 5% CO_2_ for 1h to allow binding of antibodies to the PVs. For lentiviral PVs, target cells (2 x 10^4^/well) were added and incubated for 48h at 37°C, 5% CO_2_, whereas for VSV PVs, target cells (5 x 10^4^ cells/well) were added and incubated for 24h. Then, the media was removed and discarded; Promega Bright-Glo reagent was added to the plate and incubated at room temperature for 5 min before measuring luminescence.

The data was normalized to the percentage reduction in luminescence according to the average RLU of the cell only (100% neutralization) and PV only (0% neutralization) controls and fitted into a non-linear regression model (log [inhibitor] vs. normalized response – variable slope) to interpolate the inhibitory concentrations at 50% (IC_50_)^42^.

### Generation of stable CHO cell line expressing ACE2 and TMPRSS2

A kill curve experiment was performed to establish the lowest concentration of selection antibiotics necessary to kill all cells by seeding CHO cells in 12 well plates followed by addition of antibodies starting at 0.5 µg/mL (Puromycin) and 50 µg/mL (Hygromycin). All cells died within a week with a minimum of 2 µg/mL Puromycin and 100 µg/mL of Hygromycin.

VSV-G bearing lentiviral pseudotypes were generated as described encoding lentiviral vectors MT126 pRRL-SFFV-ACE2-IRES and MT131 pRRL-SFFV-TMPRSS2.v1-IRES. CHO cells were seeded in 6-well plates (day 1) and sequencially infected with 1mL MT126 (day 2) and 1mL MT131 pseudotypes (day 3), in addition to 1 mL Ham F12. Next, cells were split and transferred to a T75 flask followed by addition of complete Ham F12 containing 2 µg/mL Puromycin and 100 µg/mL Hygromycin. After 2 or 3 passages cells were evaluated in transduction experiments using SARS-CoV-2 PVs.

### Genscript cPASS ELISA

The surrogate inhibition assay (GS-cPass; GenScript^®^, Piscataway, New Jersey, USA) was performed according to manufacturer’s instructions. Briefly, diluted samples and controls were mixed with diluted HRP-RBD solution in a 1:1 (v/v) ratio followed by 30 min incubation at 37°C to achieve 10 µg/mL of monoclonal antibody. Next, 100 µL of samples and controls were added to the ACE2 coated plate in duplicate and incubated for 15 min at 37°C. Plate was washed 4x with 1x wash solution, 100 µL TMB added to each well and incubated for 15 min in the dark. Reaction was stopped by adding 50 µL of stop solution, and plate read at 450 nm (Promega Explorer). Inhibition (%) was calculated as (1-OD sample/OD neg control) x100. The cutoff value for a positive sample is inhibition ≥ 30%.

## Results

### Lentiviral and VSV core PVs bearing SARS-CoV-2 Spike and its variants

A panel of SARS-CoV-2 PVs, including previous variants of concern, was generated bearing Spikes of ancestral (Wuhan), D614G, Alpha, Beta, Gamma, Delta and Omicron (BA.1 and BA2.75) on a lentiviral and VSV core. Additional lentiviral PV bearing Omicron variants BA.4/5 and BA.2.86 were also generated. High transduction efficiency of approximately 10^7^-10^9^ RLU/ml was achieved for all lentiviral PVs (Figure 1a), and 10^6^ RLU/ml for VSV PVs (Figure 1b) in transduction experiments of stable CHO cell lines expressing ACE2 and TMPRSS2.

**Figure 1.**
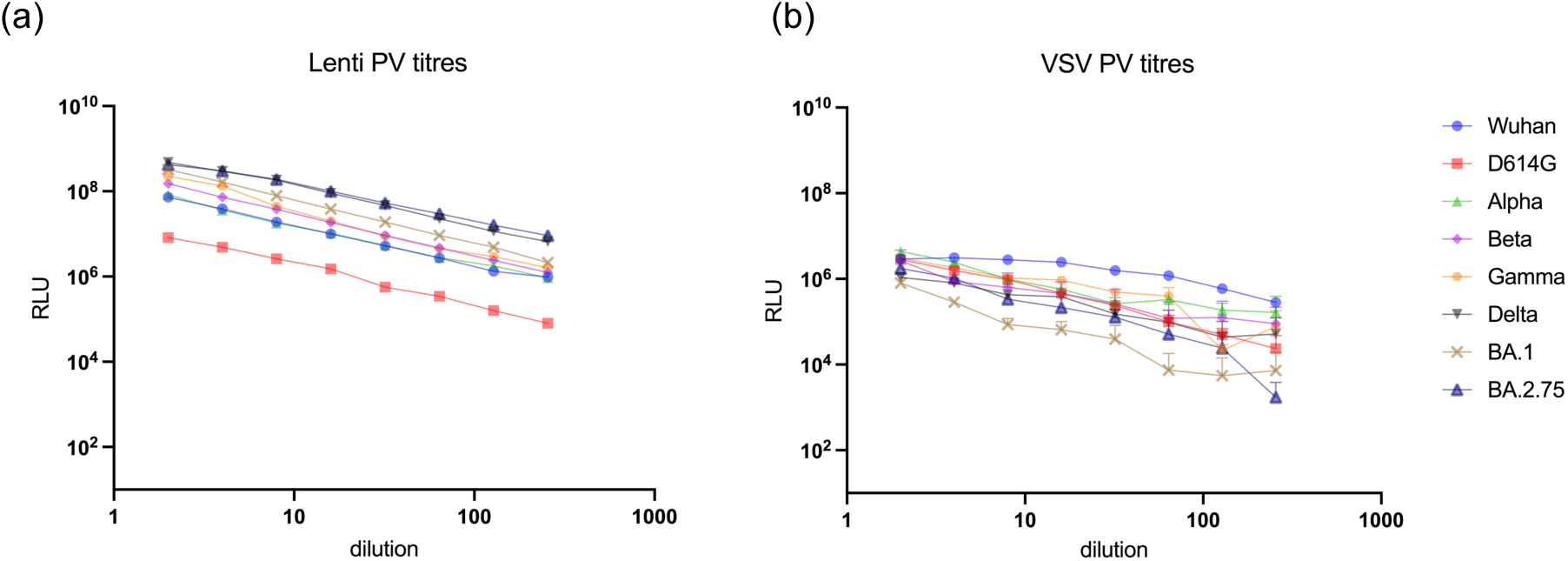
Transduction efficiency of (a) lentiviral and (b) VSV core pseudotypes bearing the main SARS-CoV-2 Spike used in this study. Each dilution point was plotted in duplicates or triplicates, expressed in relative light units (RLU) showing the mean and standard deviation (Prism v.9).

These high titre PVs yield high signal to background RLU ratio, which allows for increased accuracy in IC_50_ titre calculations. In addition, we wanted to compare both pseudotype platforms in terms of accuracy and reproducibility of results.

### Binding characteristics of monoclonal antibodies to SARS-CoV-2 antigens on Luminex

We assessed binding of a selected set of monoclonal antibodies against Wuhan, Alpha, Delta, Delta+, Eta, Lamba, Omicron (BA.1), and some mutations of importance. Most bound with similar efficiency, except to Omicron BA.1 (Figure 2, Suppl. Fig 1), where higher concentrations of mAbs were required to detect binding, except RBD mAb COV_01_242 (Figure 2). Mutations K417N and Y543F resulted in mAbs binding with stronger affinity in comparison to Wuhan, whereas mutation N439K they bound with less affinity in comparison (Figure 2). Interestingly, mAbs generated with the RBD immunogen bound to BA1 more efficiently than S1/S2 mAbs (Figure 2, Suppl. Table 1).

**Figure 2.**
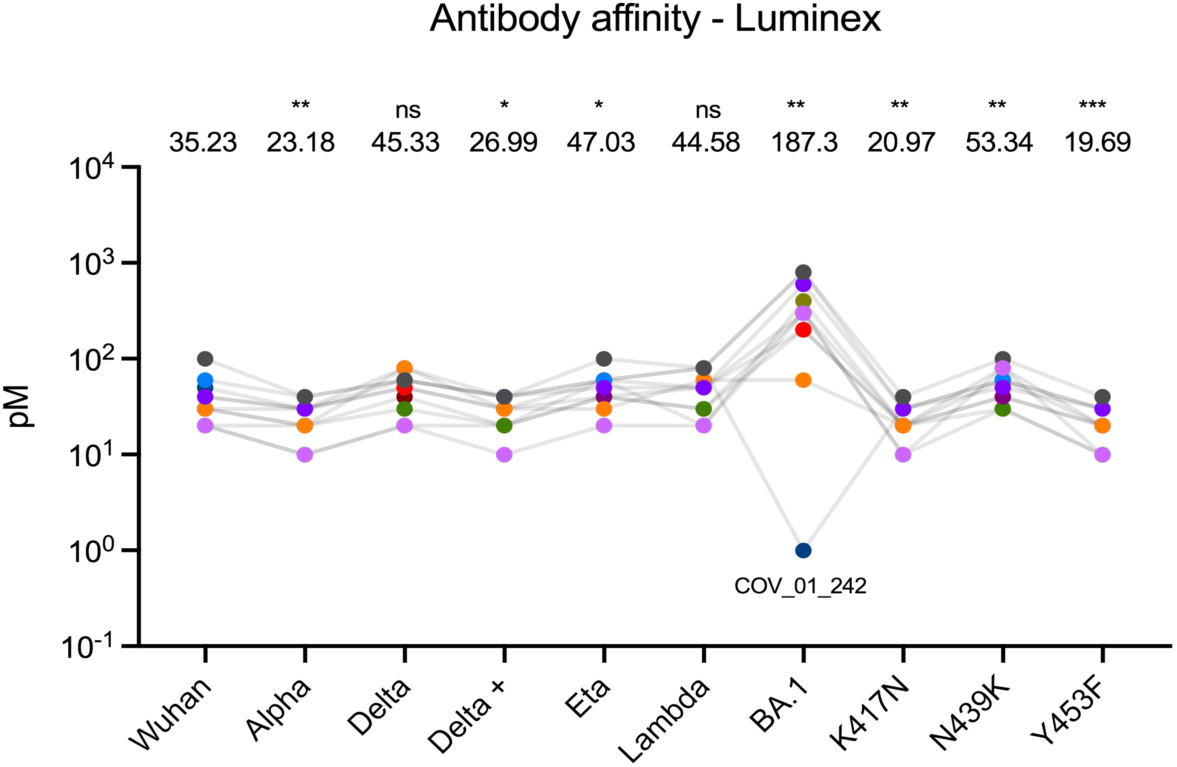
Binding characteristics by Luminex of RBD and S1/S2 monoclonal antibodies. Overall mean Kd (pM) of S1/S2 mAbs. Affinity of antibodies was calculated using the Luminex platform against antigens of selected VOC circulating at the time. The geometric mean of antibody affinity (pM) is shown above each data group. Statistical significance was determined using the Wilcoxon matched-pairs signed-rank test (Prism v.9).

### Monoclonal antibodies with strong neutralizing activity against SARS-CoV-2 VOC

Similarly, the same set of monoclonal antibodies were assessed in a neutralization assay using Luminex by addition of 600 pM of soluble ACE2-Fc as well as in a pseudotype virus neutralization assay. RBD mAb COV_01_303 did not exhibit neutralizing activity in the Luminex neutralization assay (Figure 3, Suppl. Figure 2) or in the pseudotype neutralization assay (Figure 4). Both RBD and S1/S2 mAbs neutralize Omicron BA.1 to a lesser extent (Figure 3), in agreement with the pseudotype neutralization assay (Figure 4). Overall S1/S2 mAbs retained higher neutralizing activity in comparison to RBD mAbs (Figure 3, Suppl. Table 2) against all variants tested, also in agreement with the pseudotype neutralization assay (Table 1).

**Figure 3.**
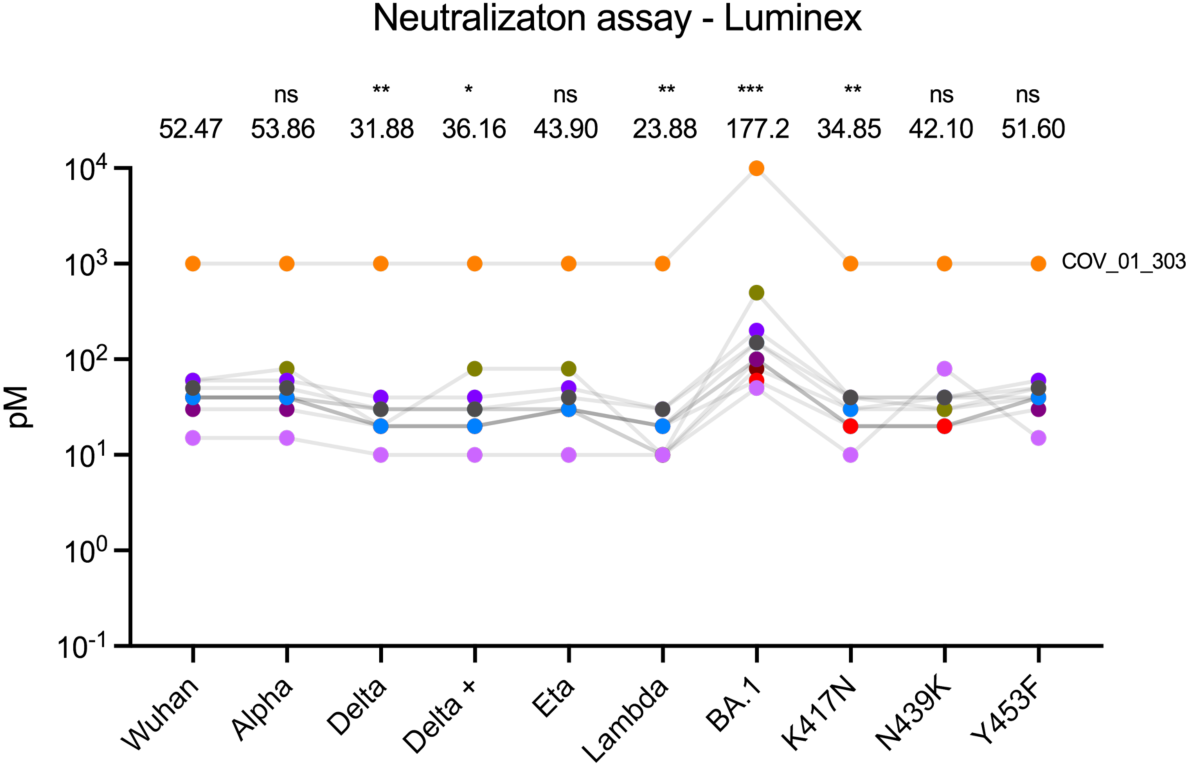
Luminex neutralization assay of RBD and S1/S2 monoclonal antibodies. The Luminex platform was employed to detect neutralizing antibodies by introducing 600 pM of soluble ACE2-Fc. The geometric mean of antibody neutralization (pM) is shown above each data group. Statistical significance was determined using the Wilcoxon matched-pairs signed-rank test (Prism v.9).

**Figure 4.**
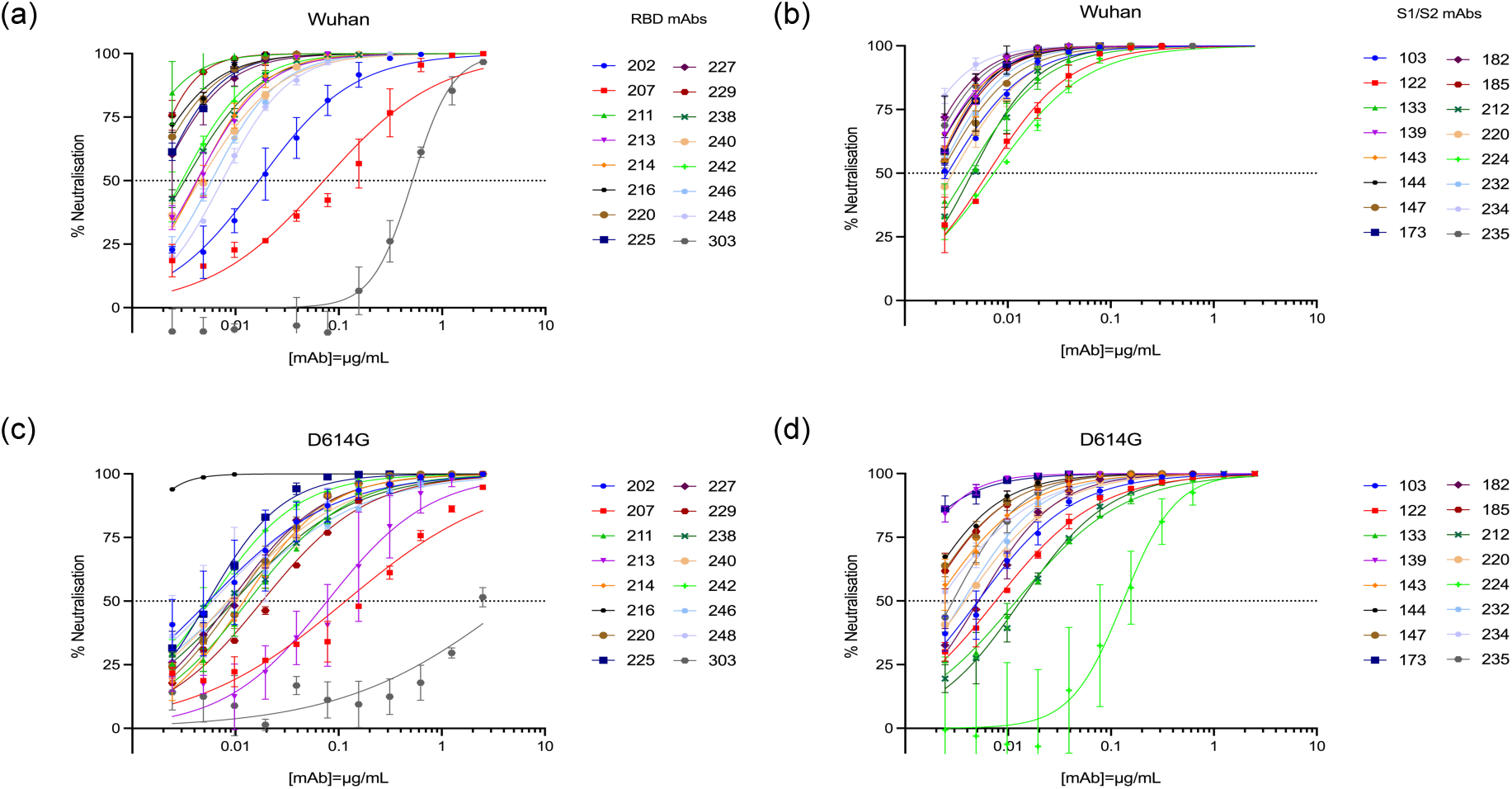
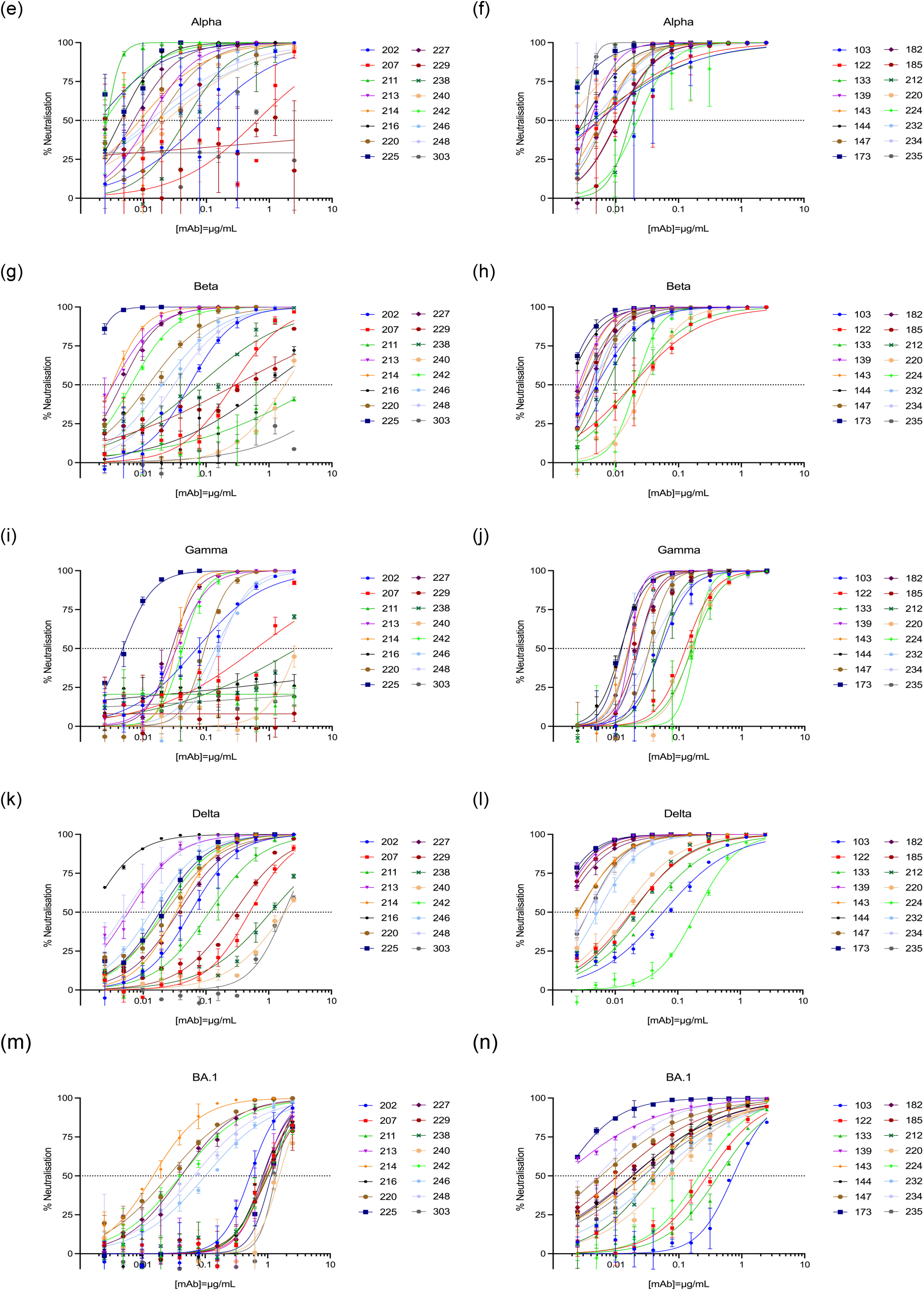
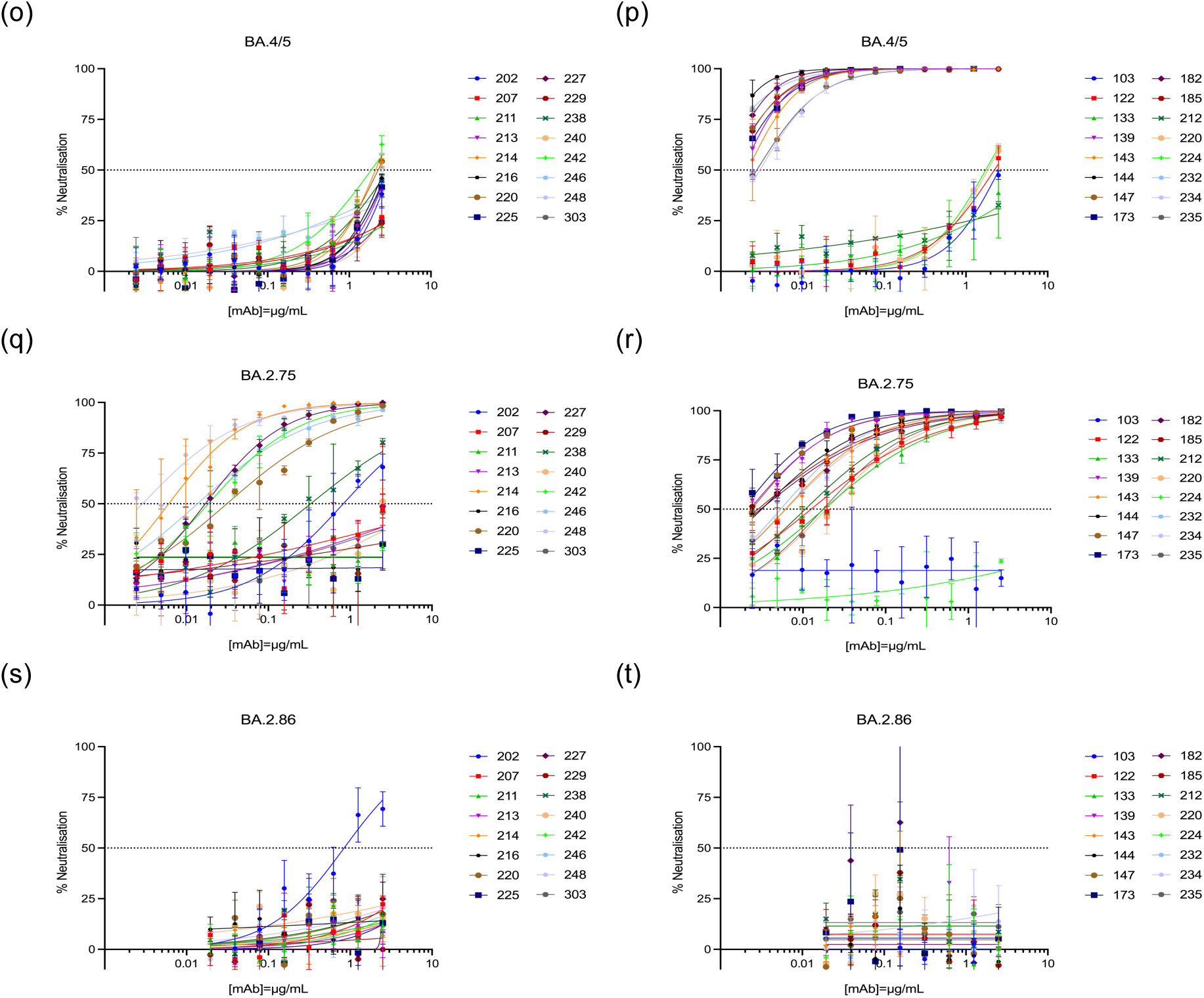
Pseudotype neutralization assay. Lentiviral pseudotypes bearing Spikes of (a,b) Wuhan, (c,d) D614G, (e,f) Alpha, (g,h) Beta, (i,j) Gamma, (k,l) Delta, (m,n) BA.1, (o,p) BA.4/5, (q,r) BA.2.75 and (s,t) BA.2.86 variants were tested against the full panel of monoclonal antibodies. Neutralization curves were grouped in mAbs raised against the RBD or S1S2 immunogen. Curves were plotted by non-linear regression (Prism v.9). Error bars indicate mean ± SD.

**Table 1.** Pseudotype neutralization assay IC_50_ against VOC. Summary table showing range of IC_50_ values across VOC: darker shade of blue (≤ 0.001 µg/mL), followed by >0.001 to ≤ 0.010 µg/mL, >0.010 to ≤0.100 µg/mL, >0.100 to ≤1 µg/mL to lighter (>1 µg/mL).

|  |  |  |  |  |  |  | 0.001 | 0.01 | 0.1 | 1 | nn |
| --- | --- | --- | --- | --- | --- | --- | --- | --- | --- | --- | --- |
|  | mAb | Wuhan | D614G | Alpha | Beta | Gamma | Delta | BA.1 | BA.4/5 | BA.2.75 | BA.2.86 |
| RBD mAbs<br>(COV_01) | 202 | 0.01688 | 0.005515 | 0.088 | 0.05502 | 0.07075 | 0.05712 | 10 | 10 | 0.8249 | 0.8417 |
|  | 207 | 0.07185 | 0.1169 | 0.6425 | 0.2697 | 0.6147 | 0.4981 | 10 | 10 | 10 | 10 |
|  | 211 | 0.0008603 | 0.01326 | 0.002637 | 6.511 | 10 | 0.1124 | 10 | 10 | 10 | 10 |
|  | 213 | 0.004231 | 0.07761 | 0.01234 | 0.00331 | 0.0336 | 0.005799 | 10 | 10 | 10 | 10 |
|  | 214 | 0.004295 | 0.01206 | 0.008715 | 0.003205 | 0.03235 | 0.03349 | 0.005568 | 10 | 0.006061 | 10 |
|  | 216 | 0.001249 | 0.0007389 | 0.004346 | 0.9725 | 10 | 0.001461 | 10 | 10 | 10 | 10 |
|  | 220 | 0.001487 | 0.009873 | 0.0172 | 0.01212 | 0.09501 | 0.02277 | 0.01656 | 10 | 0.03109 | 10 |
|  | 225 | 0.00184 | 0.005408 | 0.001864 | 0.001353 | 0.004722 | 0.01831 | 10 | 10 | 10 | 10 |
|  | 227 | 0.00183 | 0.00896 | 0.006964 | 0.004487 | 0.02843 | 0.03635 | 0.02123 | 10 | 0.01698 | 10 |
|  | 229 | 0.001392 | 0.0182 | 10 | 0.2965 | 10 | 0.2824 | 10 | 10 | 10 | 10 |
|  | 238 | 0.003238 | 0.009675 | 0.04635 | 0.07699 | 2.823 | 1.084 | 10 | 10 | 0.329 | 10 |
|  | 240 | 0.004539 | 0.008413 | 0.02291 | 1.784 | 2.798 | 1.816 | 10 | 10 | 10 | 10 |
|  | 242 | 0.003039 | 0.00567 | 0.002428 | 0.006496 | 0.04227 | 0.02042 | 0.02859 | 10 | 0.0185 | 10 |
|  | 246 | 0.005869 | 0.008978 | 0.01594 | 0.02041 | 0.1403 | 0.01341 | 0.08701 | 10 | 0.0136 | 10 |
|  | 248 | 0.007445 | 0.006018 | 0.01264 | 0.03096 | 0.1609 | 0.005087 | 0.0322 | 10 | 0.002895 | 10 |
|  | 303 | 10 | 4.978 | 10 | 10 | 10 | 1.771 | 10 | 10 | 10 | 10 |
| S1/S2 mAbs<br>(COV_08) | 103 | 0.002601 | 0.005105 | 0.005382 | 0.004975 | 0.05124 | 0.05852 | 10 | 10 | 10 | 10 |
|  | 122 | 0.006411 | 0.007574 | 0.005398 | 0.01836 | 0.1278 | 0.01817 | 0.9381 | 10 | 0.4993 | 10 |
|  | 133 | 0.004065 | 0.01203 | 0.01727 | 0.0183 | 0.1613 | 0.03223 | 3.058 | 10 | 0.5784 | 10 |
|  | 139 | 0.001628 | 0.0008523 | 0.003935 | 0.002559 | 0.0164 | 0.001249 | 0.002111 | 0.001839 | 0.02867 | 10 |
|  | 143 | 0.001935 | 0.001988 | 0.005793 | 0.002774 | 0.0157 | 0.002637 | 0.08639 | 0.002128 | 0.006162 | 10 |
|  | 144 | 0.001594 | 0.001355 | 0.003115 | 0.001719 | 0.01191 | 0.001169 | 0.03684 | 0.0008501 | 0.003175 | 10 |
|  | 147 | 0.002234 | 0.001548 | 0.006771 | 0.005024 | 0.03355 | 0.002537 | 0.005611 | 0.001226 | 0.002398 | 10 |
|  | 173 | 0.001981 | 0.0005185 | 0.001226 | 0.00162 | 0.01222 | 0.0009993 | 0.001185 | 0.00151 | 0.001882 | 10 |
|  | 182 | 0.001321 | 0.005169 | 0.01078 | 0.003018 | 0.02263 | 0.0014 | 0.04311 | 0.001143 | 0.003095 | 10 |
|  | 185 | 0.001955 | 0.001612 | 0.0105 | 0.004012 | 0.02188 | 0.00129 | 0.01954 | 0.001316 | 0.002865 | 10 |
|  | 212 | 0.004507 | 0.01322 | 0.004228 | 0.007081 | 0.04815 | 0.01894 | 0.009068 | 10 | 0.01071 | 10 |
|  | 220 | 0.002915 | 0.003817 | 0.001791 | 0.02931 | 0.1557 | 0.01297 | 0.09735 | 1.854 | 0.01924 | 10 |
|  | 224 | 0.007318 | 0.1317 | 0.02245 | 0.0207 | 0.1734 | 0.1982 | 0.1777 | 1.743 | 10 | 10 |
|  | 232 | 0.002162 | 0.003641 | 0.006649 | 0.004475 | 0.0232 | 0.005317 | 0.08509 | 0.0007175 | 0.005386 | 10 |
|  | 234 | 0.0009487 | 0.002117 | 0.004415 | 0.003924 | 0.03561 | 0.003691 | 0.007176 | 0.003006 | 0.006258 | 10 |
|  | 235 | 0.001457 | 0.002912 | 0.003267 | 0.003006 | 0.01614 | 0.003707 | 0.07416 | 0.002758 | 0.01749 | 10 |

In the pseudotype neutralization assay (pMN), many of the monoclonal antibodies exhibited strong neutralizing activity against SARS-CoV-2 and its variants (Figure 4, Table 1). Apart from mAb COV_01_303, all other mAbs neutralized PVs bearing the Wuhan Spike (Table 1). As SARS-CoV-2 continues to evolve, fewer mAbs were able to neutralize PVs, especially mAbs raised against the Wuhan RBD antigen.

(Figure 4, Table 1). Of the three different Omicron PVs tested, BA.4/5 evaded neutralization against all RBD mAbs (Figure 5a, Table 1), whereas most S1S2 mAbs neutralized BA.4/5 PVs (Figure 5b) Most mAbs raised against the full Spike retained neutralization potential (Table 1). BA.2.75 PVs were neutralized by 8 of the 16 RBD mAbs tested, whereas BA.2.86 was only neutralized by RBD mAb COV_01_202 (Figure 5a, Table 1). Overall, S1/S2 mAbs retained better neutralizing activity as the different Omicron variants emerged, except BA.2.86 (Figure 5b).

**Figure 5.**
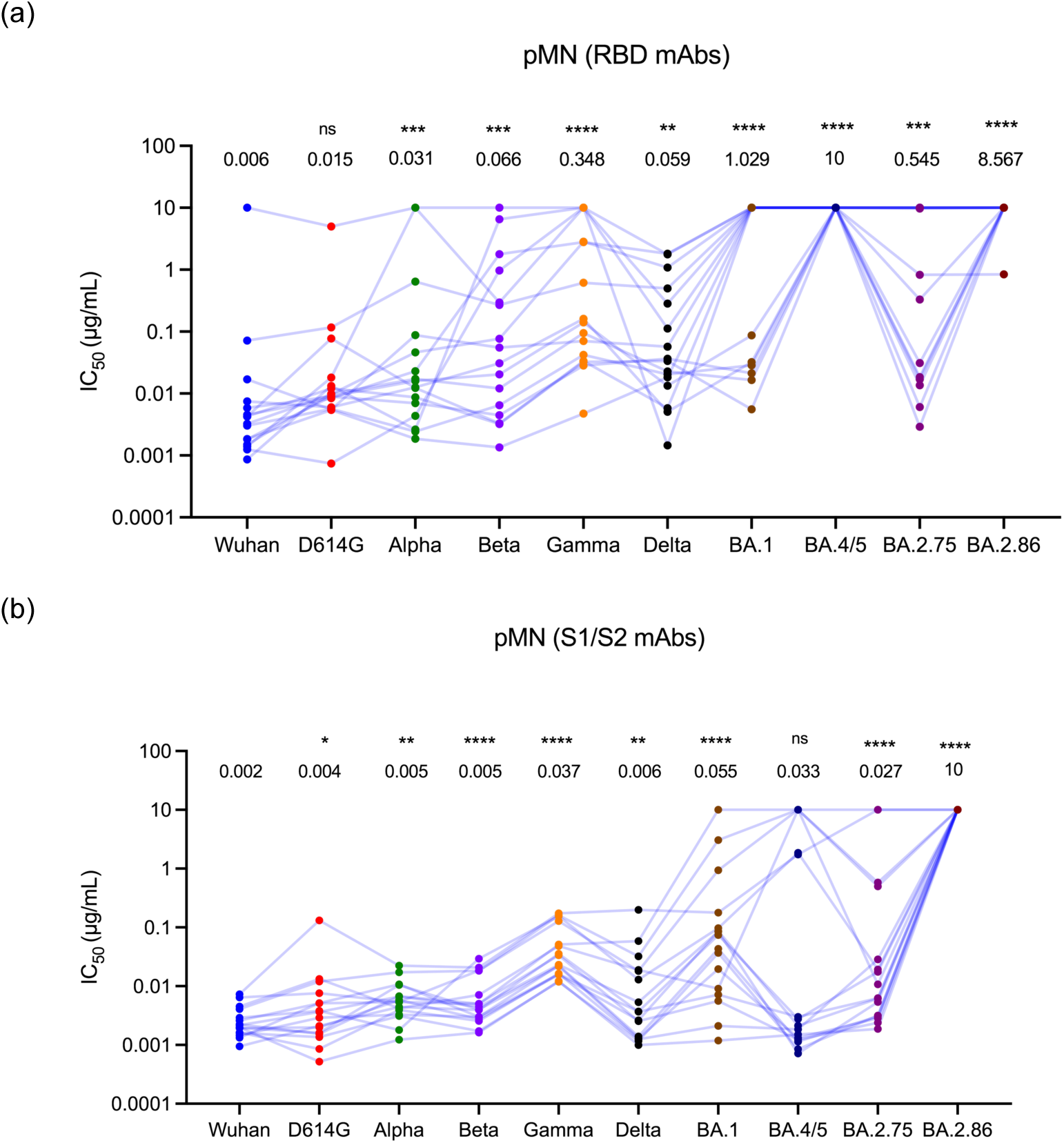
Pseudotype neutralization assay IC_50_. Results from pMN testing (a) RBD and (b) S1/S2 monoclonal antibodies. The geometric mean value of the antibody neutralization (µg/mL) is shown above each data group. Statistical significance was determined using the Wilcoxon matched-pairs signed-rank test (Prism v.9).

Gamma and the four Omicron variants tested exhibited increased evasion in neutralization against RBD mAbs (Figure 5a). Ten S1/S2 retained neutralization activity at <0.01 µg/mL against BA.4/5 PVs (Figure 5b). Overall, ten S1/S2 mAbs (COV_08_139, 143, 144, 147, 173, 182, 185, 232, 234, 235) retained neutralizing activity against most variants tested, except BA.2.86 (Table 1).

Monoclonal antibodies inhibited interaction of SARS-CoV-2 RBD with ACE2 in an ELISA-based surrogate neutralization test (Genscript cPASS)

To further asses the neutralizing activity of the panel of mAbs, the surrogate neutralization test (sVNT) Genscript cPass showed high inhibition of the RBD (Wuhan) interaction with ACE2 with all monoclonal antibodies tested at a concentration of 10 µg/mL (Figure 6), with the exception of mAb COV_01_303, which only achieved 46% inhibition (Figure 6, Suppl. Table 3). This is consistent with the results obtained from the pMN assay (Figure 4). Other variants were not available to be tested due to the fixed antigen nature of the kit.

**Figure 6:**
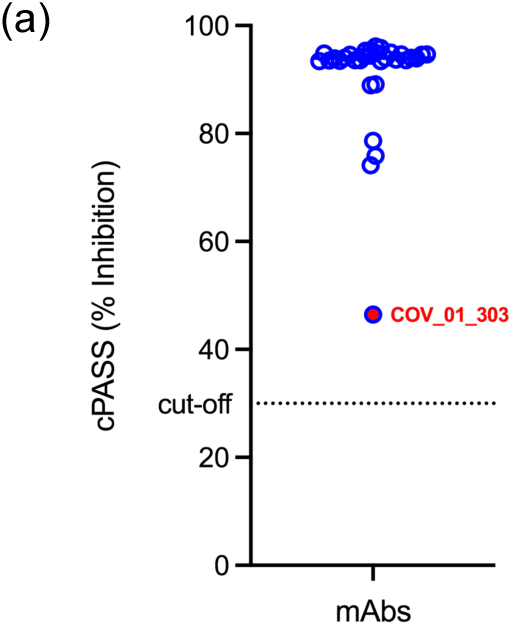
Screening of NanoTools monoclonal antibodies with the cPASS ELISA (Genscript). Results show the percentage inhibition of each monoclonal antibody. Dotted line represents the cut-off for positivity (30%). Monoclonal antibody COV_01_303 highlighted in red (Prism v.9).

### Cross reactivity against other coronaviruses and further development

The panel of monoclonal antibodies were tested against SARS-CoV-1, WIV16, RaTG13, MERS and VSV-G PVs (Figure 7, Suppl. Figure 3). RaTG3 PVs (Figure 7a) were neutralized by RBD mAbs COV_01_214 (0.079 µg/mL) and COV_01_207 (1.772 µg/mL); WIV16 PVs (Figure 7b) were neutralized by mAbs COV_01_207 (0.062 µg/mL); SARS-CoV-1 PVs (Figure 7c) were neutralized by RBD mAb COV_01_207 (1.252 µg/mL); while MERS PVs were not neutralized at all (Suppl. Figure 3).

**Figure 7.**
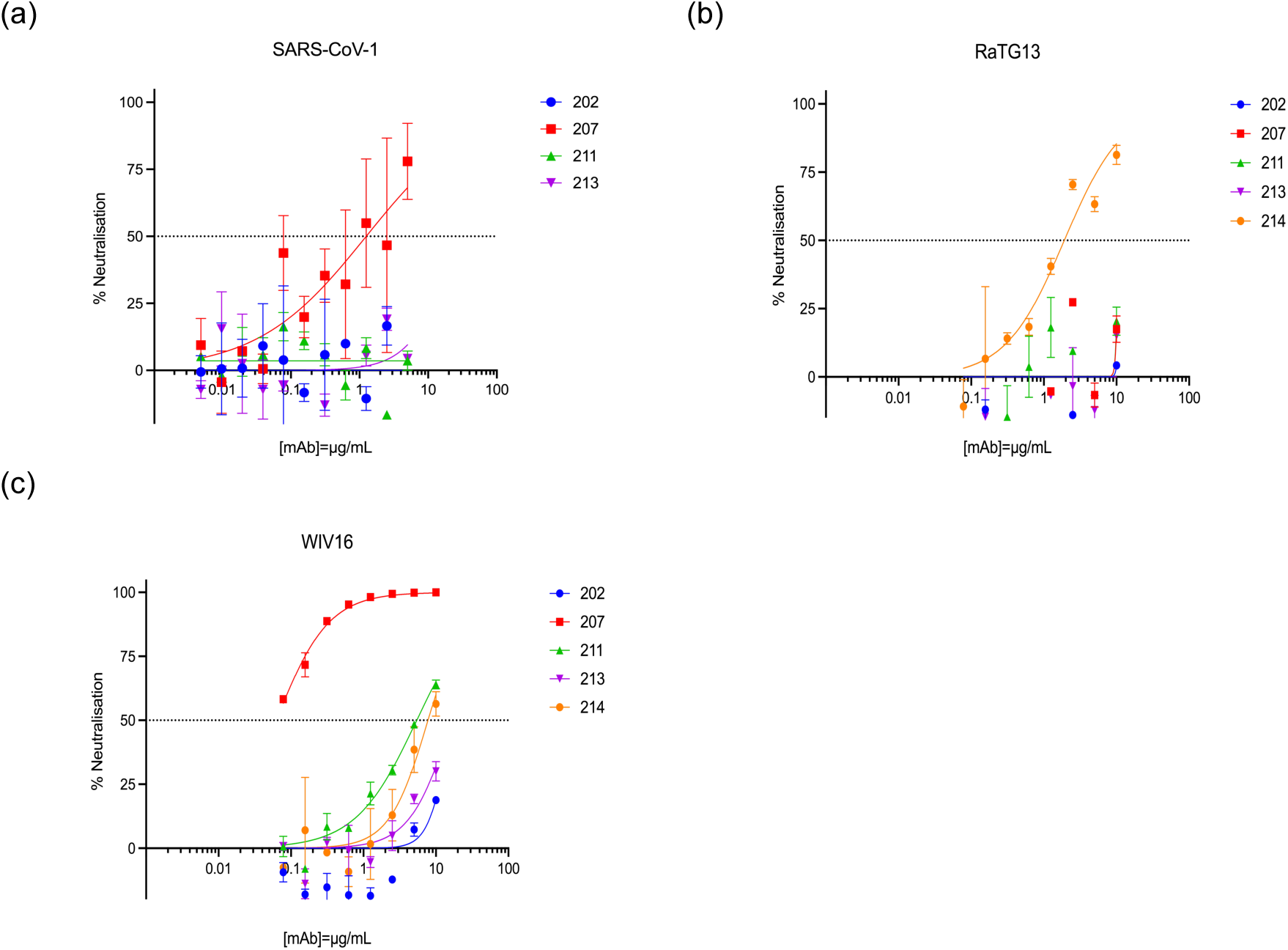
Cross reactivity with PVs (a) SARS-CoV-1 and bat viruses (b) RaTG13 and (c) WIV16. The full panel of mAbs was tested against each PV. Curves were plotted by non-linear regression (Prism v.9). Error bars indicate mean ± SD.

There was no cross-reactivity against non-CoV PVs (VSV-G) in three mAbs tested (COV_01_216, COV_08_173 and COV_08_182), as evidenced by the lack of reduction in luminescence even at higher concentrations (Suppl. Figure 3d).

Out of the 32 mAbs tested, 31 neutralized Wuhan and D614G PVs (Table 1). RBD mAb COV_01_207 neutralized most variants except BA.1 and BA.4/5 (Figure 8a,b; Table 1), as well as SARS-CoV-1 PV and bat WIV16 PV (Figure 8b). mAb COV_01_211 strongly neutralized Wuhan, D614G, Alpha (Figure 8c) and Delta PVs (Figure 8d). mAb COV_01_214 neutralized most variants (Figure 8e), except Omicron BA.4/5 and BA.2.86 (Figure 8f). It also neutralized bat virus RaTG13 (Figure 8f).

**Figure 8.**
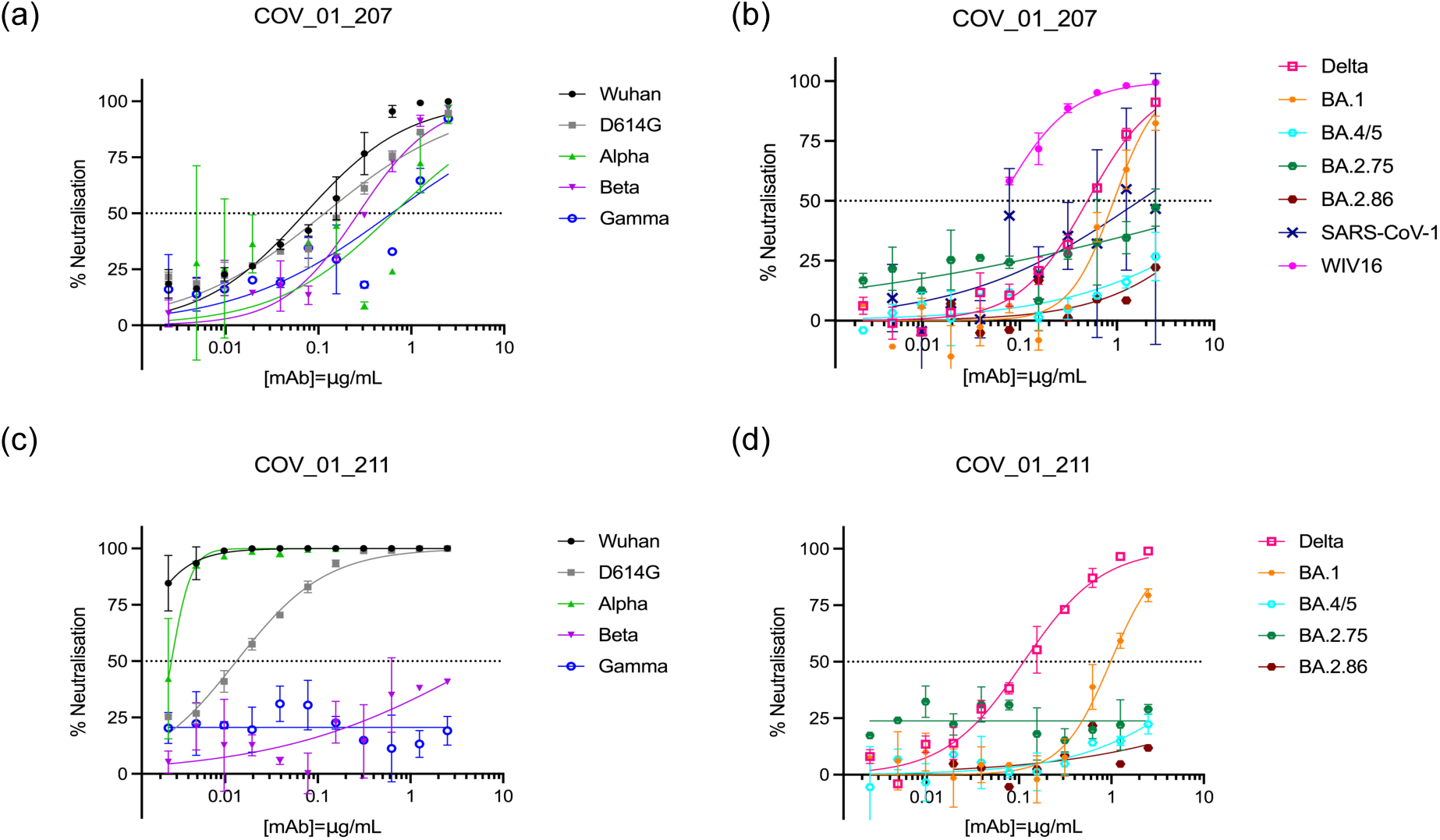
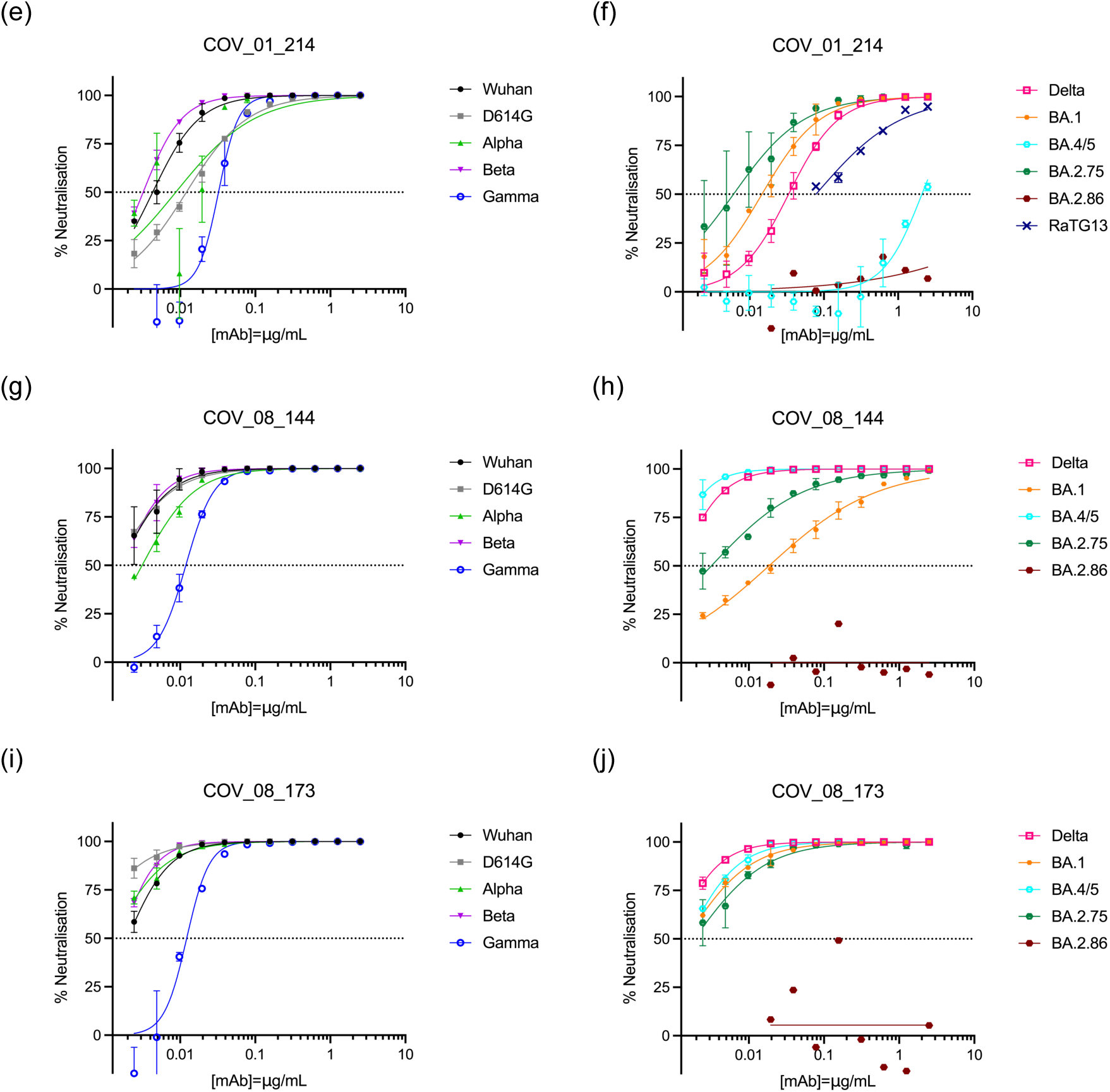
Neutralization profile of individual monoclonal antibodies (a,b) COV_01_207, (c,d) COV_01_214, (e,f) COV_08_144 and (g,h) COV_08_173. All SARS-CoV-2 variants tested are shown. Additional PVs such as SARS-CoV-1 and bat RaTG13 and WIV16 are shown where applicable. Curves were plotted by non-linear regression (Prism v.9). Error bars indicate mean ± SD.

Ten mAbs had a strong neutralization profile (Table 1), such as S1/S2 mAbs COV_08_144 (Figure 8g,h) and COV_08_173 (Figure 8i,j), which neutralized most PVs, including Omicron BA.1, BA.4/5 and BA.2.75, the only exception being BA.2.86 (Table 1).

### Comparison of lentiviral and VSV PV core platforms and stable CHO cell line

Finally, we assessed whether the lentiviral and VSV core platforms were consistent and could be used interchangeably in neutralization assays. The panel of monoclonal antibodies were tested against Wuhan (Figure 9a), Beta (Figure 9b) and Gamma (Figure 9c) lentiviral and VSV PVs.

**Figure 9.**
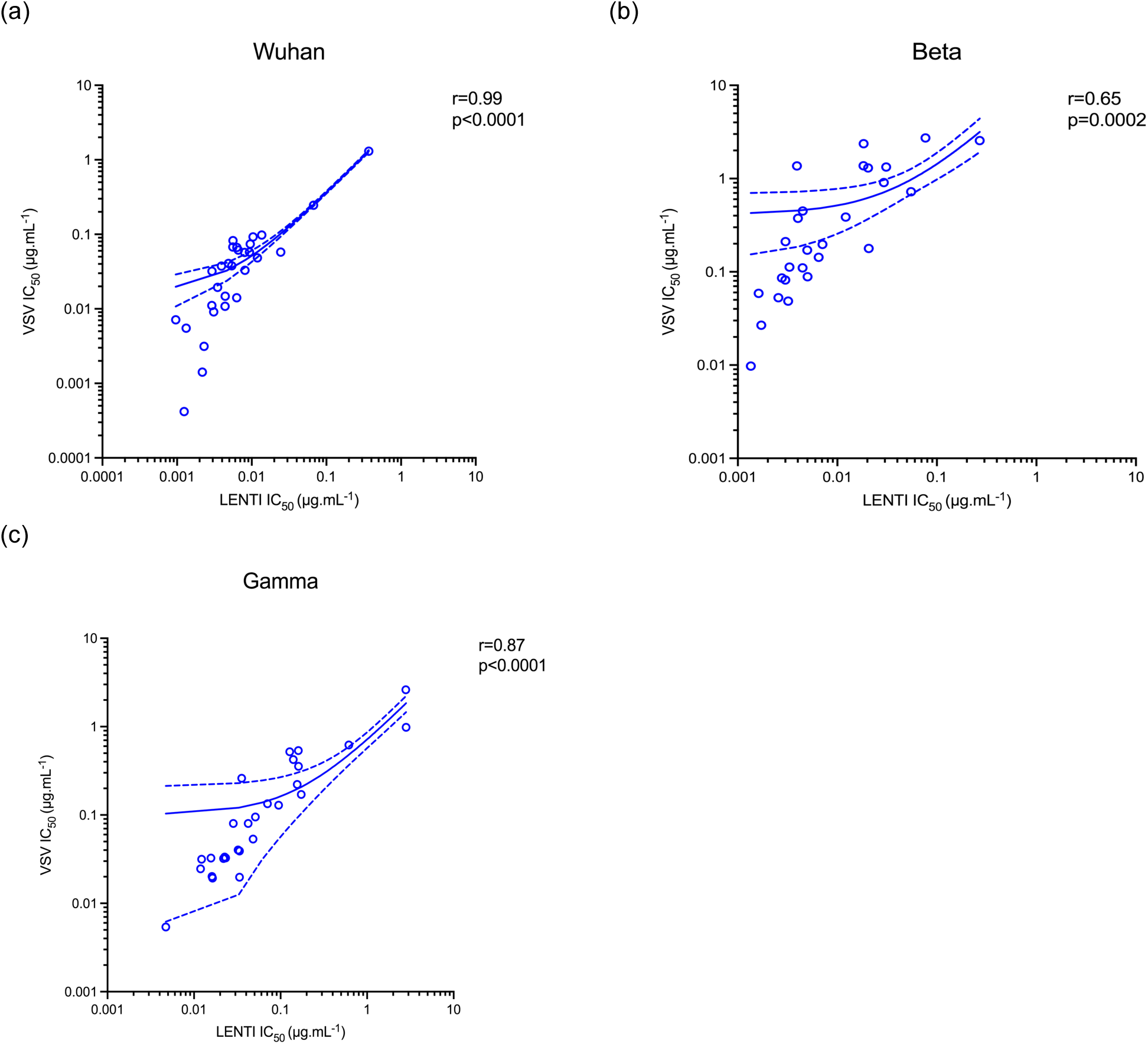
Correlation analysis between lentiviral and VSV core pseudotype neutralization assays. The panel of mAbs were tested using lentiviral and VSV core PVs bearing the Spike of (a) Wuhan, (b) Beta and (c) Gamma VOCs. The Pearson r coefficient Correlation and linear regression analyses were calculated (Prism v.9). The dashed lines are the standard deviation of the linear regression plots.

IC_50_ values were comparable: Wuhan r=0.99 (p<0.0001), Beta r=0.65 (p=0.0002) and Gamma r=0.87 (p<0.0001). However, in certain circumstances, lentiviral PVs were more sensitive than VSV PVs in neutralization assays in this study (Figure 9b).

The stable CHO cell line was evaluated in pMN (Wuhan) against pre-transfected HEK293T/17 cells with ACE2 and TMPRSS2 showing a strong correlation (r=0.99, p<0.0001) in IC_50_ values (Suppl. Figure 4).

## Discussion

The emergence of VOC, particularly the Delta variant associated with increased hospitalizations in England in comparison to the initial prevailing Alpha variant^57^ had emphasized the need for improved therapeutics against severe disease where monoclonal antibodies are being considered for use in the most severely affected patients. Even though hospitalization numbers decreased since the emergence of Omicron and its different lineages, there were still over 28,000 deaths per month worldwide in 2020^58^. In the UK, hospitalizations numbers were at 737 in the week ending 30/06/2025 (gov.uk), and according to the WHO there were 858 deaths reported worldwide (28 days to 20/07/2025). Several monoclonal antibodies have been isolated which target the Spike protein present on the viral surface, responsible for attachment to the target cell and entry after interacting with the cellular angiotensin-converting enzyme receptor (ACE2)^59–61^. Others such as tocilizumab target the IL-6 receptor reducing inflammatory responses observed in patients with cytokine storm symptoms^62,63^, was found to reduce mortality rates in patients with severe acute respiratory distress syndrome^64^.

Here we report monoclonal antibodies (mAbs) targeting the Spike of coronavirus, which strongly neutralize most VOC as well as the original Wuhan and Omicron BA.1, BA.4/5 and BA.2.75, all associated with immune escape. Mice were immunized with either the receptor binding site or the full Spike of the Wuhan original lineage, followed by isolation and purification of a panel of mAbs.

Of the 32 mAbs reported here, all bound with high affinity in the picomolar range on initial characterization by the Luminex platform (Figure 2). Most mAbs bound to Omicron BA.1 with less affinity in comparison to other VOC tested (Figure 2, Suppl. Table 1), in agreement with other reports^65,66^. Interestingly, RBD mAbs bound with higher affinity to BA.1 than S1/S2 mAbs (Figure 2, Suppl. Table 1).

NanoTools’ Luminex-based *in vitro* neutralization assay with immobilized Spike trimers and soluble ACE2-Fc enabled rapid identification of neutralizing antibodies (Figure 3). RBD mAbs (Figure 3), apart from mAb COV_01_303, neutralized most VOC tested, however BA.1 evaded neutralization (Figure 3, Suppl. Table 2). S1/S2 mAbs retained a stronger neutralization profile (Figure 3, Suppl. Table 2). This is in agreement with the pseudotype neutralization assay (Figures 4 & 5). All mAbs were screened against PV bearing the Spike of SARS-CoV-2 (Wuhan), in addition to variants of concern. IC_50_ values reached 0.001 µg/mL (Table 1). The strong neutralization profile against Wuhan was confirmed with a surrogate ELISA based assay (Genscript), where inhibition values from 74% to 96% (Figure 6, Suppl. Table 3) were observed, except for mAb COV_01_303. Previous VOCs Beta, Delta and current Omicron lineages have been associated with immune escape and increase in hospitalizations, generating concerns that monoclonal antibody therapy might be rendered ineffective^28^. Beta and Gamma contain the E484K Spike substitution thought to result in immune evasion^28,67^, therefore antibodies avoiding these sites would be beneficial. Omicron has several mutations associated with immune escape from therapeutic monoclonal antibodies^65^. In addition, mAbs targeting the N-terminal domain were found to be less effective at neutralization due to the number of mutations at this site^25^.

Omicron BA.2.86, which emerged in late summer 2023 with 30 mutations relative to XBB.1.5^68^, evaded neutralization by most mAbs tested, except RBD mAb COV_01_202 (Table 1). Mapping this epitope could be useful in identifying functional residues involved in entry as well as possible immunogens for a pan-sarbecovirus vaccine. A class III monoclonal antibody (S309) was used to identify a mutation (D339H) which caused steric hindrance and resulted in evasion by BA.2.86 and subsequent variants JN.1, Slip, FLiRT and KP.2^69^. Another mAb, iC1, was found to neutralize PVs bearing Spikes of the XBB lineage and provided protection in mice against challenge with Wuhan, BA.5 and XBB.1.5 viruses. However, it did not neutralize JN.1 PVs^70^.

We found two mAbs able to neutralize related human and bat coronaviruses. RBD mAb COV_01_207 neutralized bat virus WIV16 and human SARS-CoV-1 pseudotypes (Figure 7a) and RBD mAb COV_01_214 neutralized bat virus RaTG13 (Figure 7b). A pan-sarbecovirus neutralizing monoclonal antibody would be highly desirable as a pandemic preparedness therapeutic. We identified mAb COV_01_214 and COV_01_207 as strong mAbs that was also able to neutralize, albeit with lesser strength, RaTG13 and WIV16 respectively. We find this interesting as RaTG13 was considered to be one of the closest relatives to SARS-CoV-2, whereas WIV16 is more closely related to SARS-CoV-1, therefore it is likely that this mAb COV_01_207 is binding to a site that is evolutionarily or functionally conserved within the Spike^71^. However, any future therapeutic strategy should consider a cocktail of at least two mAbs rather than a single agent, to decrease the risk of development of resistance^72^. Updating antibody cocktails might also be necessary as other variants of concern emerge^73^. These mAbs could be good candidates to include in a future therapeutic cocktail or even playing a role in prevention of future spillovers^74^.

Similarly, heterologous antigen delivery systems might be necessary for a pan-sarbecovirus vaccine. A multiple antigen vaccine platform using ferritin with an adjuvant able to display 24 different antigens was found to be safe and well tolerated in humans^75,76^.

Neutralizing antibodies were detected over time against most SARS-CoV-2 variants using PVs, including Omicron, as well as RaTG13 and more moderately, against WIV1 and SHC014. However, no neutralizing antibodies were detected against MERS^75^. No cross-reactive antibodies against MERS were present in our panel either (Suppl, Figure 3). It is important to note that only one antigen (pre-fusion stabilized SARS-CoV-2 Wuhan Spike trimer) was present in the ferritin nanoparticle. The next step will be testing multiple antigens^75^.

When administering monoclonal antibodies in conjunction with vaccines, it is important to consider that antiviral mAbs might alter the humoral response, reducing neutralization titres due to epitope masking, therefore careful consideration to vaccination and mAb administration schedule must be given^77^.

Finally, we compared our lentiviral and VSV pseudotype platforms. We screened the panel of mAbs against Wuhan (Figure 9a), Beta (Figure 9b) and Gamma (Figure 9c) lentiviral and VSV core PVs. By calculating the Pearson correlation coefficient, all three comparisons showed a correlation between the two platforms (Figure 9). pMN with lentiviral PVs can sometimes be more sensitive than VSV PVs (Figure 9b). In our hands, one of the main advantages of lentiviral PVs is they yield higher titres (Figure 1a). However, pMNs utilizing VSV core PVs are read in 24h instead of the 48h necessary for assays with lentiviral PVs due to their cytoplasmic replication^78^, yielding quicker results. In addition, they are not affected by restriction factors such as Trim5α, which restricts infection by lentiviral PVs in certain cell lines^79^. In certain specific scenarios the VSV platform might be preferred for instance, as it has been recently found that in HIV infected individuals receiving integrase inhibitor treatment, the lentiviral platform overestimates neutralization titres^80^.

As new variants emerge, current vaccines will have to be constantly reassessed and new therapies developed. Both PV platforms are useful for vaccine evaluation. mAbs will have to be made affordable for lower income countries to be able to have an impact in the ongoing pandemic.

## Notes

### Competing Interest Statement

The authors have declared no competing interest.

