## Supplementary for "Cross-Neutralizing Monoclonal Antibodies with Broad Activity Against Human and Bat-Derived SARS-Related Coronaviruses"

### Supplementary Material:

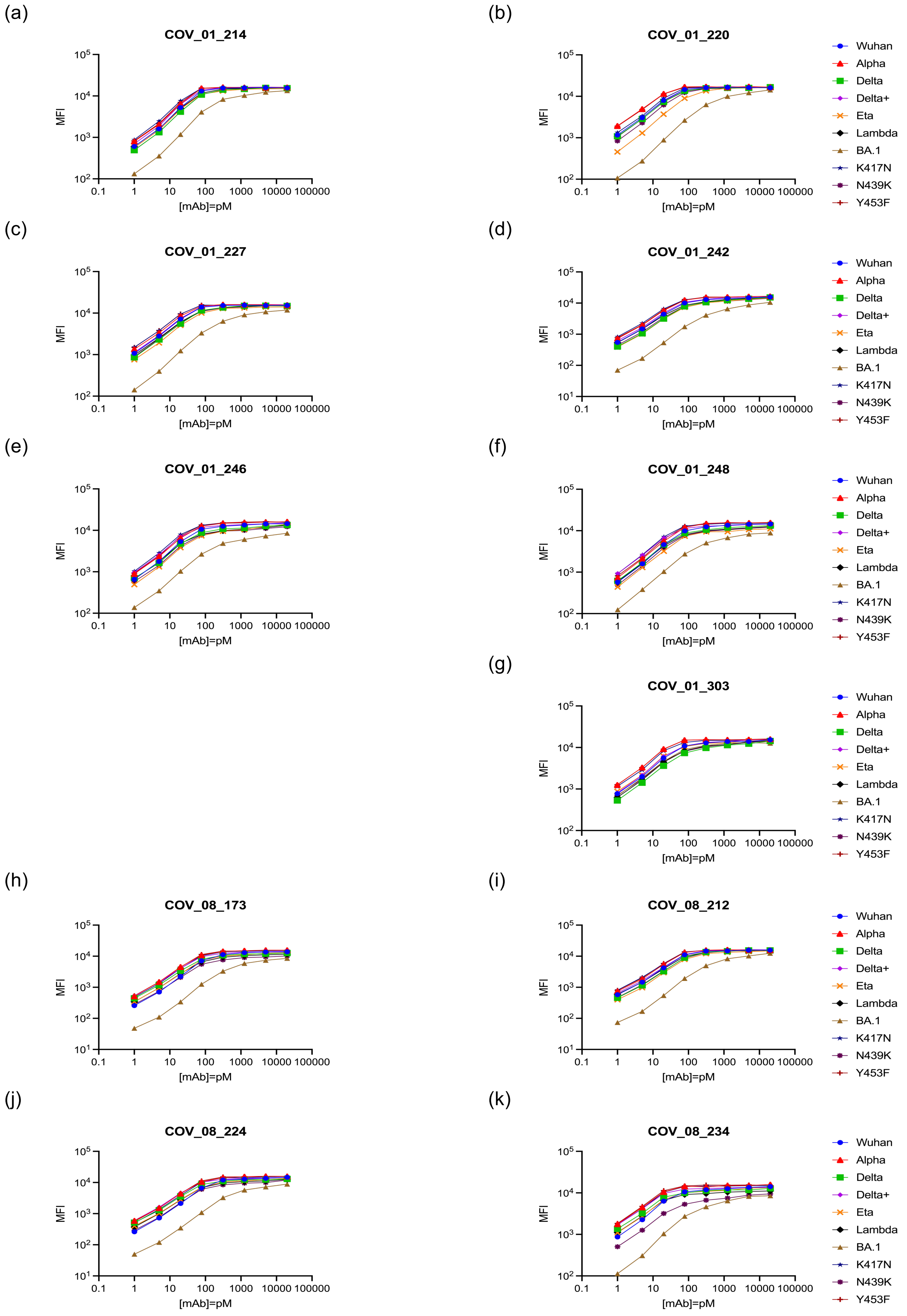

Supplementary Figure 1. Luminex assay to estimate antibody affinity. Antibodies raised from (a,b,c,d,e,f and g) RBD Wuhan and (h,i,j and k) S1S2 immunogens were tested with a selection of immobilized analytes (legend). To determine dissociation constants (Kd) binding assays were performed using a dilution series of antibodies (1-20000 pM). when fluorescent signals (MFI) are plotted against antibody concentration, the concentration at midpoint MFI corresponds to the Kd value (Prism v.9).

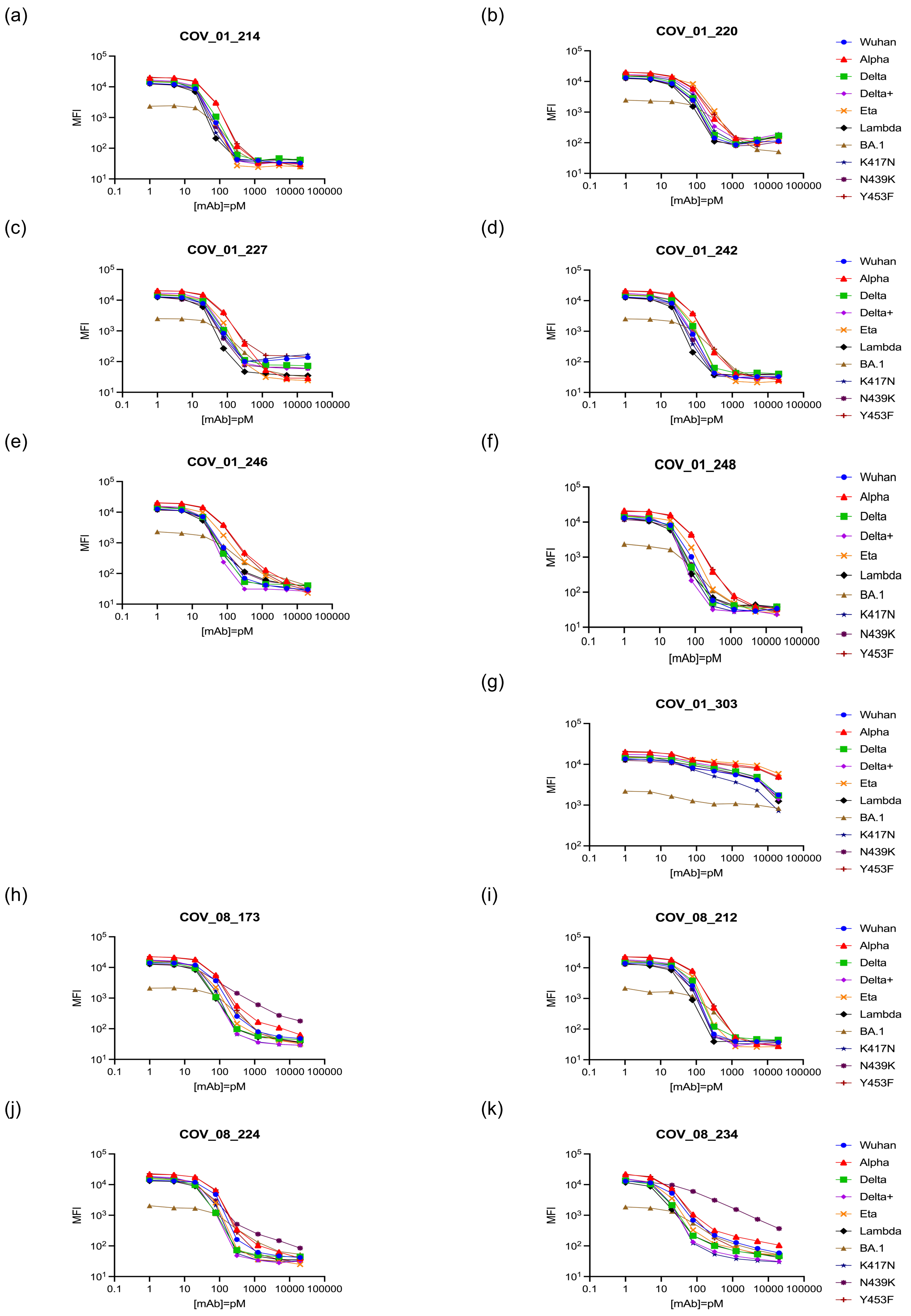
(a)
COV_01_214
(b)
COV_01_220
(c)
COV_01_227
(d)
COV_01_242
(e)
COV_01_246
(f)
COV_01_248
(g)
COV_01_303
(h)
COV_08_173
(i)
COV_08_212
(j)
COV_08_224
(k)
COV_08_234
MFI
[mAb]=pM
0.1 1 10 100 1000 10000 100000
Wuhan
Alpha
Delta
Delta+
Eta
Lambda
BA.1
K417N
N439K
Y453F

Supplementary Figure 2. Luminex assay to assess neutralization of ACE 2-Fc binding. Antibodies raised from (a,b,c,d,e,f and g) RBD Wuhan and (h,i,j and k) S1S2 immunogens were tested with a selection of immobilized analytes (legend) and a soluble analyte (600 pM ACE2-Fc). After incubation the extent of ACE2-Fc binding was determined by phycoerythrin-labelled anti-human Fc antibody. $IC_{50}$ values correspond to the antibody concentration at midpoint MFI (Prism v.9).

(a)

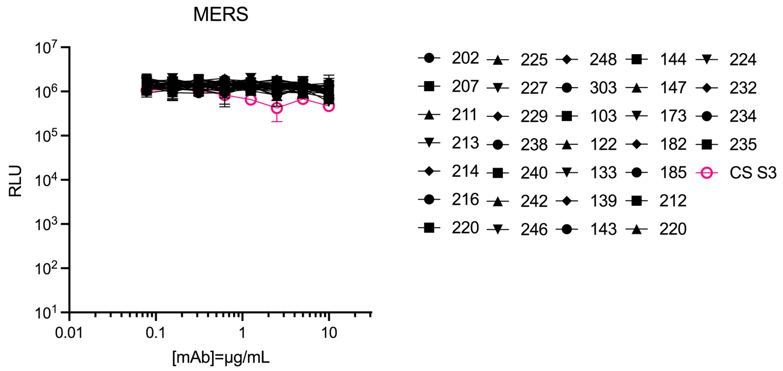


(b)

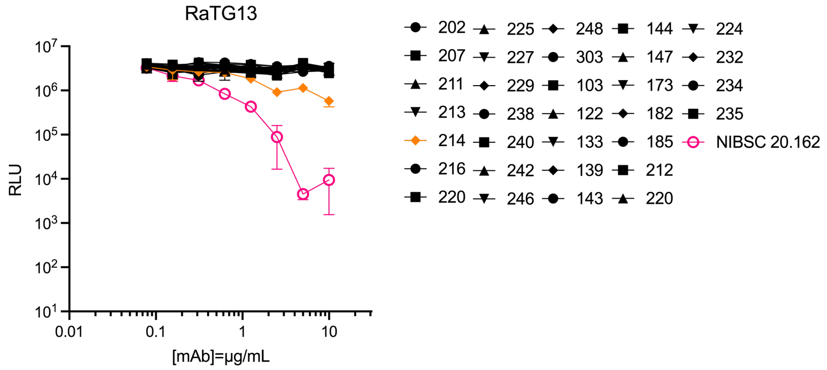


(c)

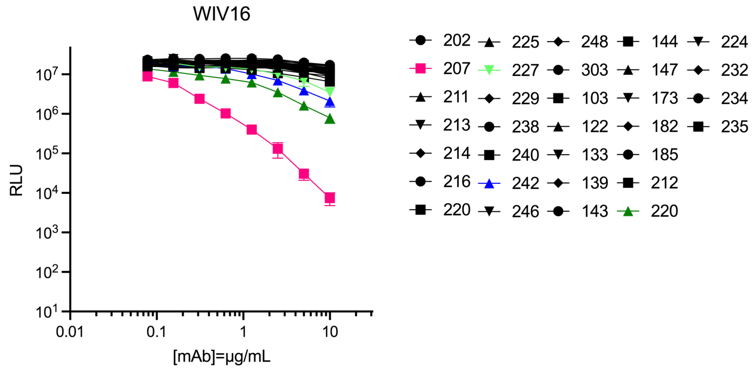


(d)

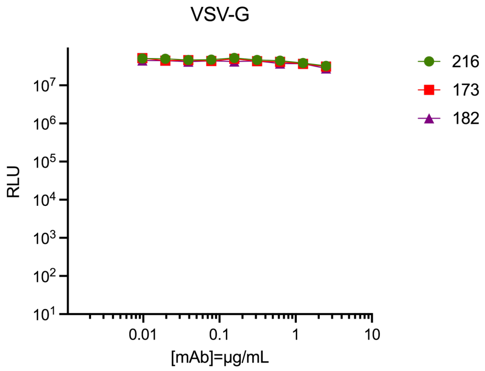


Supplementary Figure 3. Pseudotype neutralization assay with (a) MERS, (b) RaTG13, (c) WIV16 and (d) VSV-G PVs. Neutralization is evidenced by the reduction in luminescence. CS S3 anti-MERS and NIBSC Anti-SARS-CoV-2 Antibody Reagent (20/162) were used as positive controls. No neutralization was detected against MERS or VSV-G PVs (Prism v.9).

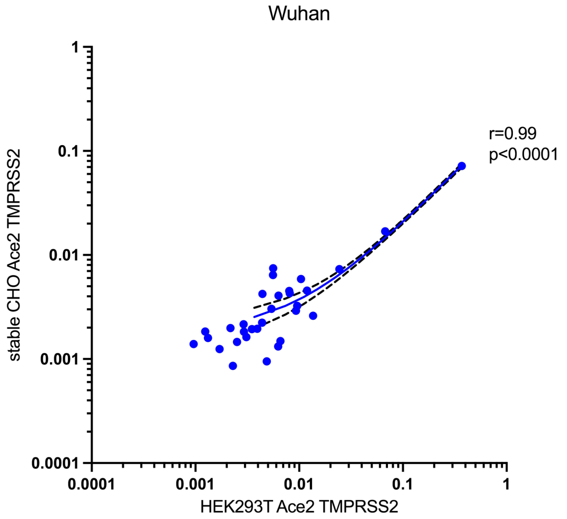


Supplementary Figure 4. Correlation analysis between a stable CHO cell line expressing ACE2 and TMPRSS2 and pre-transfected human HEK293T cells. The panel of mAbs were tested using lentiviral core PVs bearing the Spike of Wuhan. The Pearson r coefficient Correlation and linear regression analyses were calculated. The dashed lines are the standard deviation of the linear regression plots (Prism v.v9).

Supplementary Table 1. Luminex antibody affinity results (pM) from RBD (COV_01) and S1S2 (COV_08) mAbs.

| mAb | Wuhan | Alpha | Delta | Delta + | Eta | Lambda | BA.1 | K417N | N439K | Y453F |
|---|---|---|---|---|---|---|---|---|---|---|
| COV_01_214 | 30 | 20 | 40 | 20 | 40 | 30 | 200 | 20 | 40 | 20 |
| COV_01_220 | 20 | 10 | 20 | 20 | 60 | 20 | 400 | 20 | 30 | 10 |
| COV_01_227 | 20 | 20 | 30 | 20 | 40 | 30 | 300 | 10 | 30 | 10 |
| COV_01_242 | 50 | 30 | 80 | 40 | 60 | 80 | 1 | 30 | 60 | 30 |
| COV_01_246 | 30 | 30 | 50 | 30 | 40 | 50 | 300 | 20 | 40 | 20 |
| COV_01_248 | 40 | 30 | 50 | 30 | 60 | 50 | 200 | 20 | 60 | 20 |
| COV_01_303 | 30 | 20 | 80 | 30 | 30 | 60 | 60 | 20 | 80 | 20 |
| COV_08_173 | 60 | 40 | 60 | 40 | 60 | 80 | 800 | 30 | 60 | 30 |
| COV_08_212 | 40 | 30 | 60 | 40 | 50 | 50 | 600 | 30 | 50 | 30 |
| COV_08_224 | 100 | 40 | 60 | 40 | 100 | 80 | 800 | 40 | 100 | 40 |
| COV_08_234 | 20 | 10 | 20 | 10 | 20 | 20 | 300 | 10 | 80 | 10 |

Supplementary Table 2. Luminex neutralization assay results (pM) from RBD (COV_01) and S1S2 (COV_08) mAbs.

| mAb | Wuhan | Alpha | Delta | Delta + | Eta | Lambda | BA.1 | K417N | N439K | Y453F |
|---|---|---|---|---|---|---|---|---|---|---|
| COV_01_214 | 40 | 40 | 20 | 20 | 30 | 10 | 80 | 30 | 30 | 40 |
| COV_01_220 | 60 | 80 | 20 | 80 | 80 | 10 | 500 | 40 | 30 | 50 |
| COV_01_227 | 40 | 40 | 30 | 30 | 30 | 10 | 100 | 20 | 20 | 40 |
| COV_01_242 | 40 | 40 | 30 | 30 | 30 | 20 | 100 | 20 | 20 | 40 |
| COV_01_246 | 30 | 30 | 20 | 20 | 30 | 20 | 100 | 20 | 20 | 30 |
| COV_01_248 | 40 | 40 | 20 | 20 | 30 | 20 | 60 | 20 | 20 | 40 |
| COV_01_303 | 1000 | 1000 | 1000 | 1000 | 1000 | 1000 | 10000 | 1000 | 1000 | 1000 |
| COV_08_173 | 40 | 40 | 20 | 20 | 30 | 20 | 150 | 30 | 40 | 40 |
| COV_08_212 | 60 | 60 | 40 | 40 | 50 | 30 | 200 | 40 | 40 | 60 |
| COV_08_224 | 50 | 50 | 30 | 30 | 40 | 30 | 150 | 40 | 40 | 50 |
| COV_08_234 | 15 | 15 | 10 | 10 | 10 | 10 | 50 | 10 | 80 | 15 |

Supplementary Table 3. Surrogate neutralization test (cPASS ELISA, Genscript). Inhibition of the interaction of SARS-CoV-2 (Wuhan) RBD with ACE2 is considered to be any value above 30%, according to the manufacturer. The results are the average of three independent experiments.

| RBD mAb | %inhibition |
|---|---|
| 202 | 94 |
| 207 | 89 |
| 211 | 76 |
| 213 | 93 |
| 214 | 95 |
| 216 | 95 |
| 220 | 95 |
| 225 | 95 |
| 227 | 94 |
| 229 | 79 |
| 238 | 96 |
| 240 | 89 |
| 242 | 96 |
| 246 | 95 |
| 248 | 95 |
| 303 | 46 |

| S1/S2 mAb | %inhibition |
|---|---|
| 103 | 94 |
| 122 | 94 |
| 133 | 94 |
| 139 | 94 |
| 143 | 95 |
| 144 | 95 |
| 147 | 94 |
| 173 | 94 |
| 182 | 94 |
| 185 | 93 |
| 212 | 94 |
| 220 | 94 |
| 224 | 74 |
| 232 | 95 |
| 234 | 93 |
| 235 | 94 |
